# Differential GAP requirement for Cdc42-GTP polarization during proliferation and sexual reproduction

**DOI:** 10.1101/338186

**Authors:** Daniela Gallo Castro, Sophie G Martin

## Abstract

The formation of a local zone of Cdc42 GTPase activity, which governs cell polarization in many cell types, requires not only local activation but also switch-off mechanisms. Here we identify Rga3, a paralog of Rga4, as a novel Cdc42 GTPase activating protein (GAP) in the fission yeast *S. pombe*. Contrary to Rga4, Rga3 localizes with Cdc42-GTP to sites of polarity. Rga3 is dispensable for cell polarization during mitotic growth, but limits the lifetime of unstable Cdc42-GTP patches that underlie cell pairing during sexual reproduction, masking a partly compensatory patch wandering motion. In consequence, cells lacking *rga3* hyperpolarize and loose out in mating competition. Rga3 synergizes with the Cdc42 GAPs Rga4 and Rga6 to restrict Cdc42-GTP zone sizes during mitotic growth. Surprisingly, triple mutant cells, which are almost fully round, retain pheromone-dependent dynamic polarization of Cdc42-GTP, extend a polarized projection and mate. Thus, the requirement for Cdc42-GTP hydrolysis by GTPase activating proteins is distinct during polarization by intrinsic or extrinsic cues.

## INTRODUCTION

Polarization is critical for cell physiology. In most eukaryotic cell types, despite great variety of shapes and functions, small GTPases, in particular the highly conserved Rho-family GTPase Cdc42, regulate cell polarization (Etienne-Manneville, 2004). First identified for its role in the polarization of *S. cerevisiae* cells for bud formation (Adams et al., 1990), the function of Cdc42 has been conserved through evolution, as illustrated by cross-species complementation (Miller and Johnson, 1994; Munemitsu et al., 1990; Sasamura et al., 1997; Shinjo et al., 1990) and its requirement for polarization in numerous cell types, including the fission yeast *S. pombe* (Miller and Johnson, 1994), *C. elegans* oocytes (Gotta et al., 2001; Kay and Hunter, 2001), Drosophila neuroblasts (Atwood et al., 2007) or mammalian epithelia and oocytes (Wang et al., 2013; Wu et al., 2007).

Cdc42 is under complex regulation and cycles between active and inactive states (Vetter and Wittinghofer, 2001). When bound to GTP, Cdc42 activates effectors, including nucleators of actin assembly, such as formins, regulators of vesicle secretion, such as the exocyst complex, and PAK-family kinases (Perez and Rincon, 2010). These collectively convert a localized Cdc42 signal into effective cell polarization. Cdc42 activation relies on guanine nucleotide exchange factors (GEFs), which promote exchange of GDP for GTP. For its inactivation, Cdc42 has intrinsic GTPase activity, which is also promoted by GTPase-activating proteins (GAPs). Cdc42, which associates with membranes through a prenyl moiety, can also be sequestered in the cytosol by GDP dissociation inhibitors (GDIs) (DerMardirossian and Bokoch, 2005). Importantly, cycling of Cdc42 GTP-bound, active and GDP-bound, inactive states is critical for its function in cell polarization. In fission yeast, both Cdc42 disruption and constitutive activation lead to cell rounding and lethality, with disruption causing small round cells and constitutive activation large ones (Bendezu et al., 2015; Miller and Johnson, 1994). In consequence, the local activity of Cdc42 is critical for cell polarization.

Local activity results in part from localized GEFs, of which there are two in *S. pombe*, Scd1 and Gef1. These are together essential and localize at sites of growth and division (Coll et al., 2003; Hirota et al., 2003). However, local activation is not sufficient to generate locally restricted Cdc42 activity: cells also need to inactivate Cdc42 to prevent the spread of the active form. For instance, we recently observed that the cell pole-restricted activity of the related small GTPase Ras1 requires inactivation by its GAP, the deletion of which leads to loss of spatial information and Ras1-GTP distribution over the whole plasma membrane (Merlini et al., 2018). In fission yeast, two Cdc42 GAPs, Rga4 and Rga6, both of which localize to cell sides, contribute to the spatial restriction of active Cdc42 to the cell tips, with *rga4*Δ and *rga6*Δ mutant cells exhibiting enlarged cell width (Revilla-Guarinos et al., 2016; Tatebe et al., 2008). However, even double mutant cells retain polarized Cdc42-GTP zones, albeit a bit wider, suggesting that negative controls of Cdc42 activity remain in place. Cdc42 inactivation may also involve detachment from the membrane and sequestration in the cytosol by GDI. In *S. cerevisiae*, the GDI plays an important role in promoting Cdc42-GDP recycling to the polarity patch and accumulation of Cdc42 at its site of activity, with GAP activity also recently implicated (Slaughter et al., 2009; Woods et al., 2016). In *S. pombe*, Cdc42-GDP exhibits fast lateral mobility at the plasma membrane, such that the slower-diffusing Cdc42-GTP form can polarize independently of the GDI (Bendezu et al., 2015). Consistently, deletion of the sole GDI yields no overt phenotype (Bendezu et al., 2015; Nakano et al., 2003).

Cell polarization can take many shapes and forms for which the same polarity components are re-used. In fission yeast, Cdc42 activity can respond to intracellular microtubule-deposited landmarks that recruit it to cell poles (Kokkoris et al., 2014; Tatebe et al., 2008). There, it oscillates in an anti-correlated manner (Das et al., 2012) to drive the bipolar growth patterns of these cells and defines cellular dimensions during mitotic growth (Bendezu et al., 2015; Kelly and Nurse, 2011). Upon spore germination, Cdc42-GTP polarizes spontaneously, forming dynamic zones that stabilize upon spore outgrowth (Bonazzi et al., 2014). During sexual differentiation, Cdc42-GTP also forms unstable zones that explore the cell periphery upon pheromone exposure and become stabilized upon increased pheromone concentrations (Bendezu and Martin, 2013). In the physiological process, cells of opposite mating type (P and M) express and secrete distinct pheromones and cognate receptors, which activate a common Ras1-MAPK signalling cascade (Merlini et al., 2013). In potential partner cells, Cdc42 patches act as a source of pheromone, such that facing patches in partner cells stabilize each other, driving cell pairing (Merlini et al., 2016). Consequently alterations in Cdc42 patch dynamics modify partner cell choice. For instance, excessive pheromone perception compromises Cdc42 dynamics outside the cell tip region, leading to preferential mating between just-divided sister cells (Bendezu and Martin, 2013); excessive Ras1 activity promotes zone stabilization at reduced pheromone concentration, leading to defective cell pairing (Merlini et al., 2016).

The oscillatory patterns described above imply the existence of positive and negative feedbacks. Positive feedbacks are well described from work in *S. cerevisiae*, where they underlie the spontaneous polarization of Cdc42 in absence of landmarks (Gulli et al., 2000; Irazoqui et al., 2003; Wedlich-Soldner and Li, 2003). One prominent feedback involves the formation of a complex between a Cdc42 effector PAK kinase and a Cdc42 GEF through a scaffold protein, thought to amplify initial stochastic variations in Cdc42 activity to promote symmetry-breaking (Kozubowski et al., 2008). The molecular controls of negative feedback that destabilize the patch remain largely unclear. Phosphorylation of the Cdc42 scaffold by the PAK kinase may contribute in dampening the positive feedback reaction (Das et al., 2012; Howell et al., 2012; Rapali et al., 2017). Mechanical constraints from the cell wall may contribute to patch destabilization (Bonazzi et al., 2014). The role of Cdc42 GAP proteins in promoting negative feedback has not been systematically explored.

Here, we describe a novel Cdc42 GAP, Rga3. We show that Rga3 is a paralog of Rga4, which arose from gene duplication in the *Schizosaccharomyces* lineage. In contrast to Rga4 and Rga6, Rga3 is recruited to sites of Cdc42 activity, yet it synergizes with these two GAPs during mitotic growth to restrict Cdc42-GTP zone size and cell dimensions. During pheromone-dependent polarization, Rga3 is recruited to the Cdc42 patch where it promotes its dynamics and modulates partner choice. Surprisingly, a triple GAP mutant, though lacking polarity during mitotic growth, retains almost complete ability to polarize during sexual differentiation and mate, indicating fundamental differences in Cdc42-GTP zone regulation in distinct contexts.

## RESULTS

### Rga3 is a paralog of Rga4

Two Cdc42 GAPs have been reported in *S. pombe:* Rga4 and Rga6 (Revilla-Guarinos et al., 2016; Tatebe et al., 2008). Recent work showed that Rga6 collaborates with Rga4 in the control of cell dimensions, as double deletion of *rga4* and *rga6* leads to shorter and wider cells than either *rga4Δ* or *rga6Δ* single mutants (Revilla-Guarinos et al., 2016). However, the phenotype of this double mutant is much weaker than that caused by overexpression of a constitutively active allele of Cdc42 (Cdc42^Q61L^), which leads to complete polarity loss and formation of round cells with cytokinesis defects (Fig. 1A-B) (Bendezu et al., 2015; Miller and Johnson, 1994). This discrepancy suggests the existence of other GAP(s) promoting Cdc42-GTP hydrolysis.

**Figure 1:**
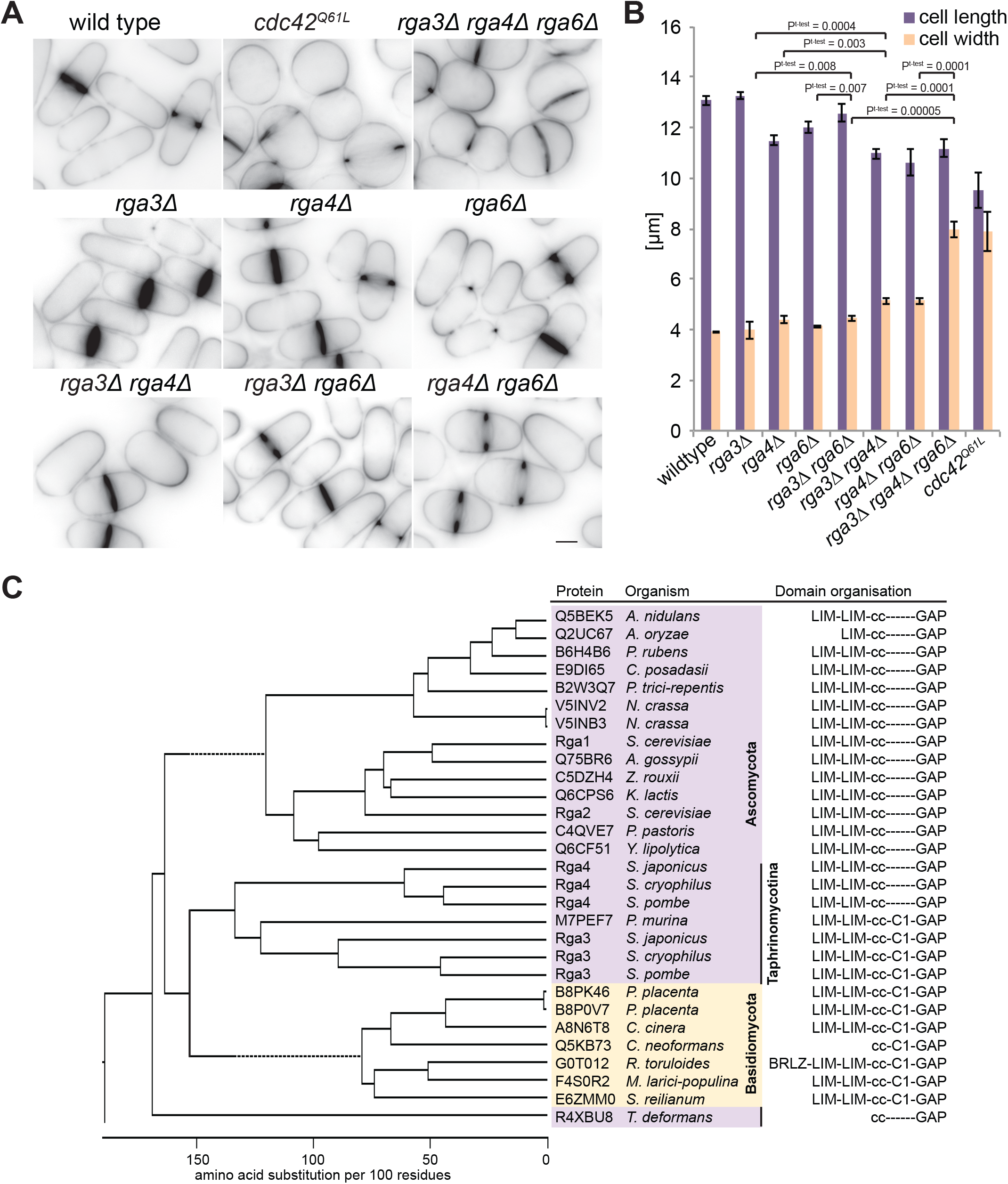
Rga3 is a paralog of Rga4 and contributes to cellular dimensions. (A) Medial plane inverted images of indicated strains stained with calcofluor. Bar = 3μm. (B) Average cell length and width of the indicated strains grown in EMM-ALU at 25°C (N = 3 experiments, with n>145 cells each). Error bars show the standard deviation. (C) Phylogenetic tree of ClustalW alignment of selected RhoGAP domain-containing proteins from Ascomycetes (purple) and Basidiomycetes (yellow) species. The Taphrinomycotina group to which fission yeasts belong is highlighted. The domain architecture of each protein (derived from SMART analysis) is shown on the right. Note that the C1 domain is present in basidiomycetes and some taphrinomycotina sequences, but absent in all other ascomycete sequences.

BLAST searches with Rga4 against the fission yeast genome revealed Rga3 as reciprocal best hit. Further searches for RhoGAP domain-containing sequences in ascomycetes and basidiomycetes, and phylogenetic analysis of proteins from selected species, showed that Rga3 and Rga4 cluster together on the same branch as the *S. cerevisiae* Cdc42 GAPs Rga1 and Rga2 (Fig. 1C). Reciprocal BLAST analysis did not reveal any clear orthologue pair with the *S. cerevisiae* GAPs, consistent with the notion that *S. cerevisiae* underwent a whole-genome duplication during its evolution (Wolfe and Shields, 1997). Furthermore, with the exception of *S. cerevisiae* and a few other species, all ascomycetes and basidiomycetes contain a single *rga4-* like gene, whereas all *Schizosaccharomyces* species have two (Fig. 1C). Thus, Rga3 and Rga4 likely arose from gene duplication in the *Schizosaccharomyces* lineage.

All Rga4-like GAPs share the same domain structures: two N-terminal LIM domains, a central coiled-coil region, and a C-terminal RhoGAP domain (Fig. 1C). Interestingly, Rga3 contains an additional C1 domain, predicted to bind lipids (Kaibuchi et al., 1989; Ono et al., 1989). This domain architecture is not shared by any other ascomycete protein except by the basal *Taphrinomycotina* group, to which the *Schizosaccharomyces* lineage belongs. Rga4-like GAPs also exhibit similar domain architecture, including a C1 domain, in the closely related basidiomycete fungi. BLAST searches with Rga3 or Rga4 GAP domain through other eukaryotic phyla returned Chimaerin-family RacGAPs as closest hits, which also bear a linked C1 domain (Canagarajah et al., 2004). Thus, a likely scenario is that the Rga3/4 ancestral state contained a C1 domain, which was subsequently lost in Rga4 and other ascomycete species.

### Rga3 is a Cdc42 GAP

Deletion of *rga3* did not lead to any evident morphological phenotype during vegetative growth. Indeed, cells lacking *rga3* showed similar cell length and width as wildtype cells (Fig. 1A-B). However, deletion of *rga3* in combination with *rga4Δ* and *rga6*Δ led to cells significantly wider and more rounded compared to the single or double mutants, a phenotype very similar to that of overexpression of a constitutively active allele of Cdc42 (Fig. 1A-B). Similarly, *rga3*Δ *rga4*Δ and *rga3*Δ *rga6*Δ double mutants were also significantly wider than either single mutant. These phenotypes strongly suggest that Rga3 could be a Cdc42 GAP.

The dramatic morphologic change in *rga3Δ rga4Δ rga6Δ* cells raised the question of how Cdc42 activity is distributed along the cortex. To address this question, we co-imaged Cdc42-mCherry^SW^, a functional internally mCherry-tagged allele of Cdc42 encoded at the endogenous locus (Bendezu et al., 2015), and CRIB-GFP in wildtype and *rga3Δ rga4Δ rga6Δ* strains. The CRIB domain, which specifically binds Cdc42-GTP, is used as a marker for active Cdc42. In contrast to the restricted distribution of CRIB around the tips of wildtype cells, CRIB formed very large cortical domains occupying broad zones all over the cell cortex in the triple mutant, with Cdc42-mCherry^SW^ showing a similar distribution (Fig. 2A). Cortical profile measurements confirmed a much broader distribution of both Cdc42 and its active form (Fig. 2B). Of note, the very tight overlap between the normalized Cdc42 and CRIB distribution profiles, indicative of Cdc42 protein enrichment to sites of activity in wildtype cells (Bendezu et al., 2015), was largely preserved in the triple mutant condition. Thus, the correlation between Cdc42 activity and its local enrichment does not depend on GAP proteins (Fig. 2A). The Cdc42-GTP-binding scaffold protein Scd2 (Endo et al., 2003; Wheatley and Rittinger, 2005) similarly localized to a broad cortical region (Fig. 2C). We observed a broad variation of zone sizes for both CRIB and Scd2, ranging from zones covering an enlarged cell tip, to almost complete cell surface coverage. These results indicate that Rga3 in conjunction with other GAP proteins serve to restrict the size of the Cdc42-GTP domain.

**Figure 2:**
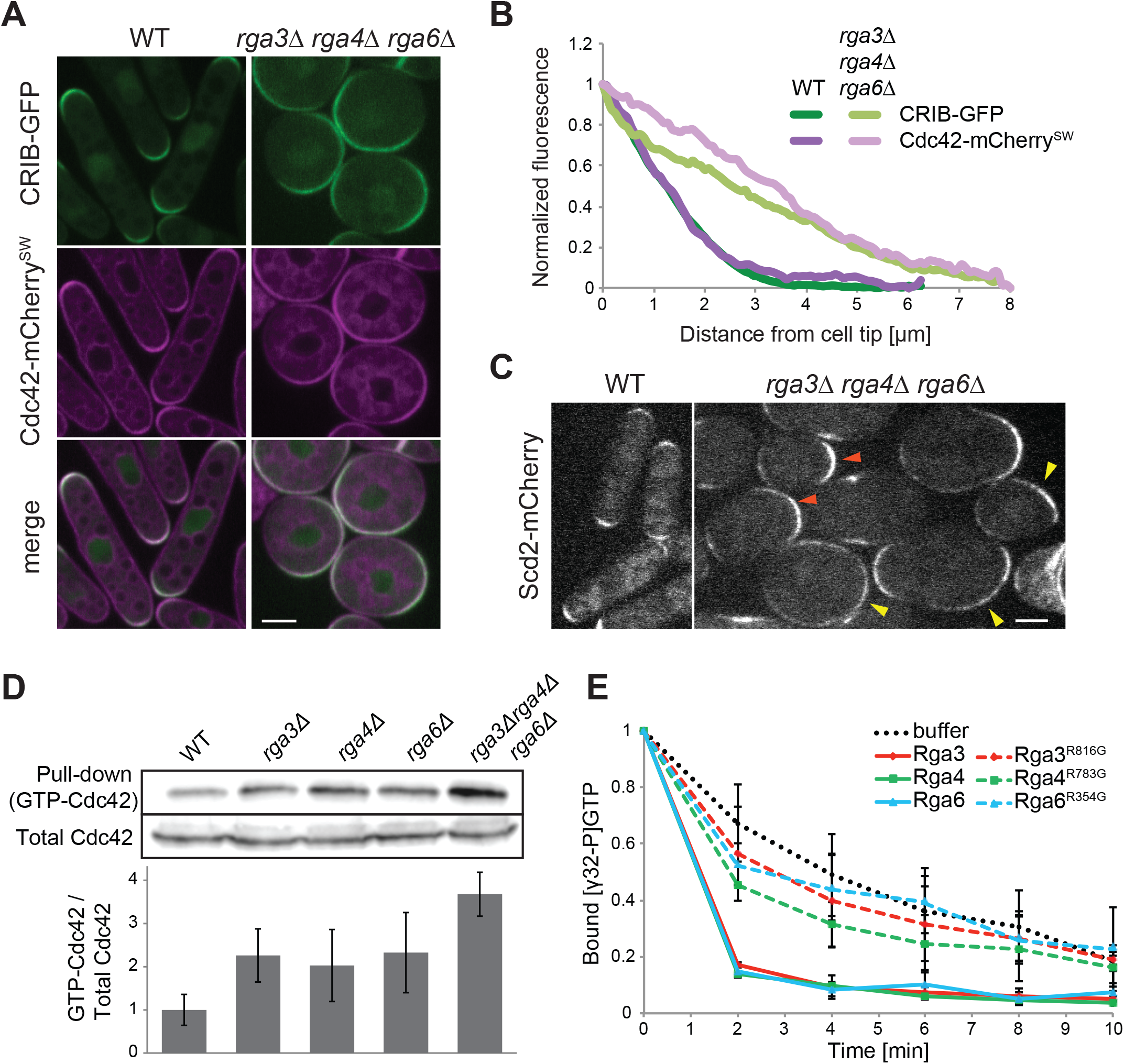
Rga3 is a Cdc42 GAP. (A) Medial plane spinning disk confocal images of Cdc42-mcherry^SW^ and CRIB-GFP in WT and *rga3Δrga4Δrga6Δ*. (B) Average cortical profiles of Cdc42 and CRIB fluorescence intensity normalized to the maximum and minimum intensity values. For *rga3Δrga4Δrga6Δ* profiles, only cells in which CRIB-GFP formed defined zones were used (n ≥ 20), so the described phenotype is underestimated in this quantification. (C) Medial plane spinning disk confocal images of Scd2-mcherry in WT and *rga3Δrga4Δrga6Δ*. Orange arrowheads indicate enlarged zones at cell poles; yellow arrowheads point to zones extending over a large part of the cell cortex. (D) Cdc42-GTP pull-down assay. GST-CRIB was coupled to glutathione beads and mixed with extracts of wild type, *rga3Δ, rga4Δ, rga6Δ* or *rga3Δrga4Δrga6Δ* cells expressing Cdc42-mCherry^SW^. Cdc42 was revealed in input and pull-down fractions with anti-mCherry antibody. One representative experiment is shown on top, the average of three independent experiments is reported on the graph below. (E) *In vitro* GAP assay. [γ-^32^P]GTP-loaded recombinant Cdc42 was incubated with recombinant full length GAPs that were wildtype or catalytically inactive (point mutations indicated) or with buffer and Cdc42-associated radioactivity was determined by scintillation counting. The data was normalized to the value before GAP addition. Bar = 3μm. Error bars show standard deviation.

To test whether Rga3 is indeed a Cdc42 GAP, protein extracts from wildtype, single *rga3*Δ, *rga4*Δ *and rga6*Δ mutants and triple mutants carrying Cdc42-mcherry^SW^ were used to determine the total amount of GTP-bound Cdc42 by pull-down with glutathione S-transferase (GST)–CRIB. Extracts of cells lacking *rga3* showed an increase in GTP-bound Cdc42 compared to wildtype extracts, comparable to the levels shown upon *rga4* and *rga6* deletion. Moreover, the triple mutant presented a further increase in the fraction of active Cdc42 (Fig. 2D), consistent with the additive phenotypes seen in vivo.

To further verify Rga3’s direct role as a Cdc42 GAP, we performed an *in vitro* GTPase assay on Cdc42. Purified recombinant Cdc42 pre-loaded with [γ-32P]GTP was incubated with recombinant Rga3, Rga4 or Rga6 to test the ability of these GAPs to increase the rate of Cdc42-GTP hydrolysis. All three GAPs, including Rga3 increased the rate of Cdc42-GTP hydrolysis. Point mutations in the catalytic arginine finger of each GAPs, predicted to affect GTP hydrolysis (Scheffzek et al., 1998), blocked their ability to increase the intrinsic GTPase activity of Cdc42 (Fig. 2E). Taken together, these data demonstrate that Rga3 is a Cdc42 GAP.

### Rga3 colocalizes with active Cdc42 independently of its GAP domain and its cortical localization is mainly dependent on its C1 domain

Given the observation that Rga3 and Rga4 are likely paralogs, have similar protein structure, and share function in the regulation of Cdc42, we compared the localization of the two GAP proteins. Tagging Rga3 with GFP yielded a functional protein as judged by the morphology of *rga4*Δ *rga6*Δ *rga3-GFP* cells, which was indistinguishable from that of *rga4*Δ *rga6*Δ cells (see Fig 3G). Strikingly, while Rga4 forms clusters at cell sides (Das et al., 2007; Tatebe et al., 2008), Rga3 localized to sites of cell division and cell tips, where its distribution was similar to that of Cdc42 (Fig. 3A-B). Rga3 was also cytosolic. Furthermore, whereas Rga4-RFP covers the non-growing cell pole of monopolar *tea1*Δ cells (Tatebe et al., 2008), Rga3-GFP was present only at the growing cell pole (Fig. S1A). Thus, Rga3 localization to cell tips correlates with active growth.

**Figure 3:**
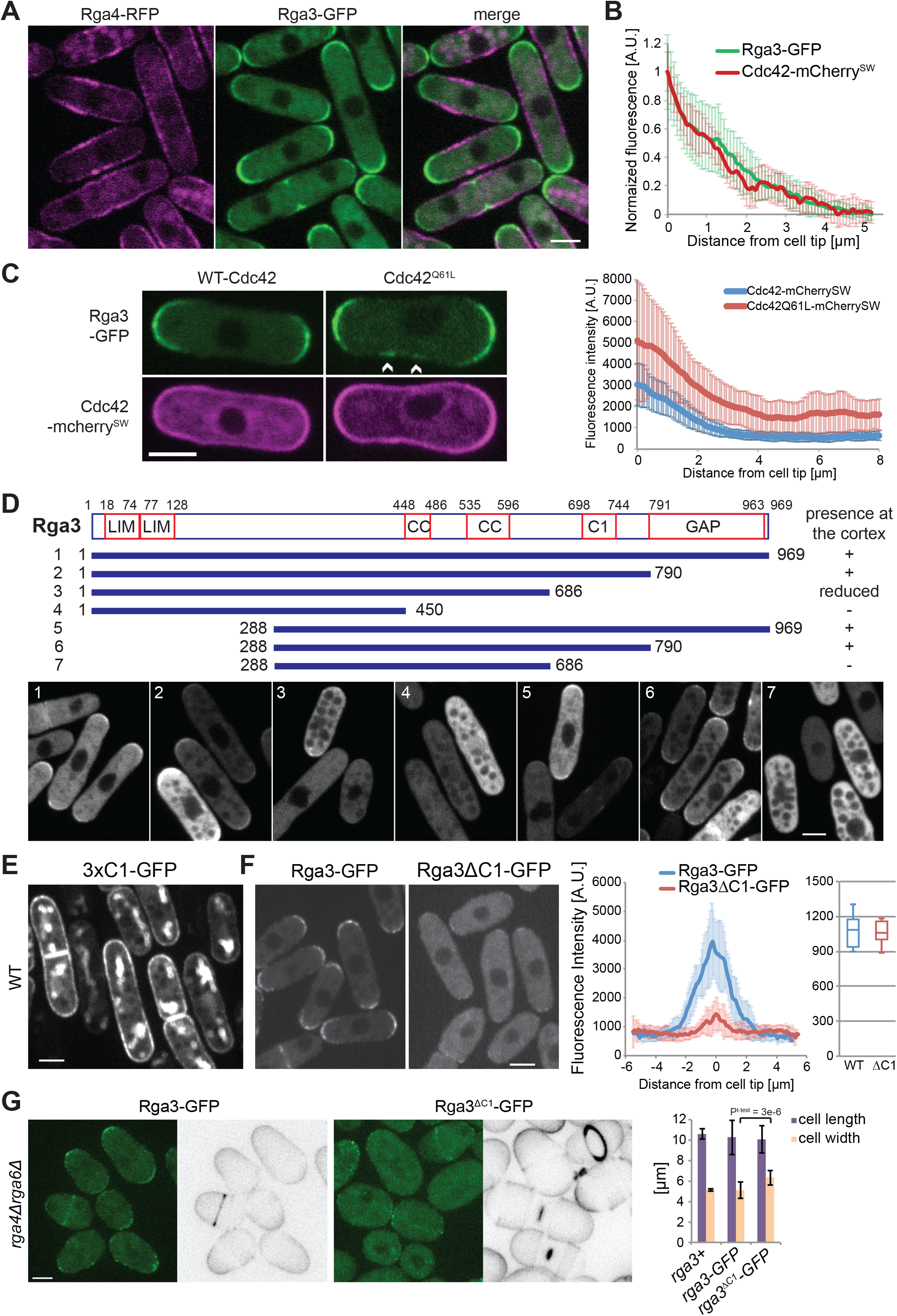
Rga3 colocalizes with active Cdc42 and binds the cortex through its C1 domain. (A) Images of Rga3-GFP and Rga4-RFP. (B) Average cortical profiles of Cdc42-mcherry^SW^ and Rga3-GFP fluorescence intensity normalized to the maximum and minimum intensity values. n=24 tips. (C) Images of Rga3-GFP strains expressing a pREP41-Cdc42-mcherry^SW^ (WT or Q61L) plasmid (left). Arrowheads indicate zones of Rga3-GFP presence at the cell sides. Average cortical profiles of Rga3-GFP fluorescence (right). In the strain carrying the Cdc42^Q61L^ allele, only rod-shaped cells were analyzed. n=18. (D) Schematic representation of Rga3 fragments whose localization was tested upon GFP-tagging and plasmid expression. Cortex-binding ability is summarized on the right. Representative images of *rga3Δ* cells expressing the plasmids are shown at the bottom. (E) Image of cells expressing a triple tandem copy of Rga3 C1 domain from plasmid. (F) Images of Rga3-GFP and Rga3^ΔC1^-GFP expressed from the native genomic locus (left). Average cortical profiles of Rga3-GFP and Rga3^ΔC1^-GFP fluorescence intensity (middle). Whole cell intensity of Rga3-GFP and Rga3^ΔC1^-GFP (right). n=11. (G) Calcofluor and GFP fluorescence images of *rga4Δrga6Δ* cells expressing Rga3-GFP or Rga3^ΔC1^-GFP (left). Average cell length and width of the same strains (n > 50) (right). All micrographs are medial plane spinning disk confocal images. Bars = 3μm. Error bars show standard deviation.

Rga3 cell tip localization was largely independent of F-actin, vesicular trafficking and microtubules: treatment with Latrunculin A (an actin depolymerizing agent), Brefeldin A (a drug that blocks vesicular trafficking) or Methyl Benzimidazol-2-yl-Carbamate (MBC, a microtubule depolymerizing agent) did not lead to delocalization of Rga3 from the tips of the cell (Fig. S1B). By contrast, we found that Rga3 localization was dependent on Cdc42-GTP. Indeed, cells expressing from plasmid a constitutively GTP-bound allele of Cdc42 (Cdc42^Q61L^), which decorates the entire cell cortex, showed Rga3 localization at the sides of the cells in addition to cell tips (Fig. 3C). Note that Cdc42^Q61L^ was expressed here for a shorter amount of time than in Fig. 1A to visualize Rga3 localization in rod-shaped cells before cell rounding. This suggests that Rga3 is directly or indirectly recruited to cell poles by active Cdc42.

To further understand the determinants of Rga3 localization, we generated truncated GFP-tagged forms of Rga3, expressed from plasmids, and assessed their localization in wildtype and *rga3*Δ cells (Fig. 3D). The full-length protein localized at cell tips, as expected, and no difference was observed between wildtype and *rga3*Δ backgrounds (Fig. 3D, fragment 1). Interestingly, truncation of the GAP domain did not impair Rga3 localization to the cortex, suggesting recruitment to cell poles does not occur through direct interaction between Cdc42-GTP and the GAP domain (Fig. 3D, fragment 2). By contrast, further truncation of the C1 domain, or the N-terminus containing both LIM domains, reduced, and in combination abolished, Rga3 cortical localization (Fig. 3D, fragments 3, 5, 6 and 7). The central coiled-coil regions also contributed, as truncation of these in combination with the C1 domain abrogated cortical localization (Fig. 3D, fragment 4). We conclude that multiple Rga3 domains contribute to its localization at cell tips, with the C1 domain playing a critical role. Of note, all fragments that associated with the cell cortex were enriched at the cell tips, not the cell sides, indicating that the multiple determinants of Rga3’s cell tip recruitment do not mask an Rga4-like cell side localization.

Because the C1 domain is essential for Rga3 localization in the absence of the LIM or coiled-coil domains and is unique to Rga3, we further probed its function. A construct with three tandem copies of the C1 domain decorated the entire cell cortex, consistent with the predicted role of this domain as lipid binder (Fig. 3E). We constructed an Rga3 allele lacking only the C1 domain (Rga3^ΔC1^-GFP) expressed as sole copy from the native genomic locus. Rga3-GFP and Rga3^ΔC1^-GFP were expressed at similar levels, but with distinct localization pattern: while Rga3-GFP decorated the whole tip cortex, Rga3^ΔC1^-GFP was mainly cytosolic with only a few punctae of weak fluorescence intensity at the cell tips (Fig. 3F). The C1 domain was also required for Rga3 function, as the deletion of the C1 domain in an *rga4Δ rga6Δ* background produced rounded cells, similar to *rga3Δ rga4Δ rga6Δ* triple mutants (Fig. 3G). We conclude that the C1 domain of Rga3, though not sufficient to direct Rga3 to cell poles, is a major contributor of its localization to sites of polarity.

Because Rga3 is indirectly recruited by active Cdc42, it may form a negative feedback on Cdc42 activity. Cdc42 activity has been shown to oscillate between the two cell tips in an anticorrelated manner, a behavior that requires negative feedback (Das et al., 2012). We thus compared Cdc42 oscillations in wildtype and *rga3Δ* cells. However, analysis of CRIB-GFP signal oscillations over time showed a similar anti-correlation between the two cell poles in the two strains, with a small decrease in frequency in *rga3Δ* cells (Fig. S2). Thus, consistent with the lack of morphological phenotype of *rga3Δ* cells, these observations indicate that negative regulation by Rga3 only functions redundantly with other Cdc42 GAPs during the vegetative life stage.

### Rga3 limits growth projection in response to pheromone and confers a competitive advantage during sexual reproduction

The absence of detectable phenotype of *rga3Δ* cells raises the question of what selective pressures promoted the maintenance of the *rga3* gene after gene duplication. We found that in contrast to the lack of morphological phenotype during vegetative growth, *rga3*Δ cells exhibited alterations in mating morphologies, extending long or poorly oriented shmoos. In a first series of experiments, *h*+ WT or *rga3*Δ cells expressing Scd2-GFP were mixed with wildtype *h-* cells and allowed to mate. Almost 30% of *h+ rga3Δ scd2-GFP* cells were found to extend a projection yet fail to find a partner after 20h in mating conditions (Fig. 4A-B). When observed in timelapse experiments, these cells did not always persistently grow from a single pole, as 10% alternated between the two poles (Fig. 4A, Movie S1). Pair-forming *rga3Δ* cells also extended longer shmoos and consequently formed pairs with more distant cells: while the majority of wildtype cell pairs formed between cells initially distant by <1.5μm, most *rga3Δ* x wt pairs formed between cells distant by >2μm (Fig. 4A, 4C). In a second set of experiments, we treated h-cells lacking the P-factor protease Sxa2 with increasing concentrations of synthetic P-factor pheromone and quantified the proportion of shmooing cells after 24h. *rga3Δ* cells formed shmoos at lower pheromone concentrations than *rga3+* cells (Fig. 4D), suggesting a higher sensitivity to pheromone consistent with the aberrant formation of shmoos in mating mixtures.

**Figure 4:**
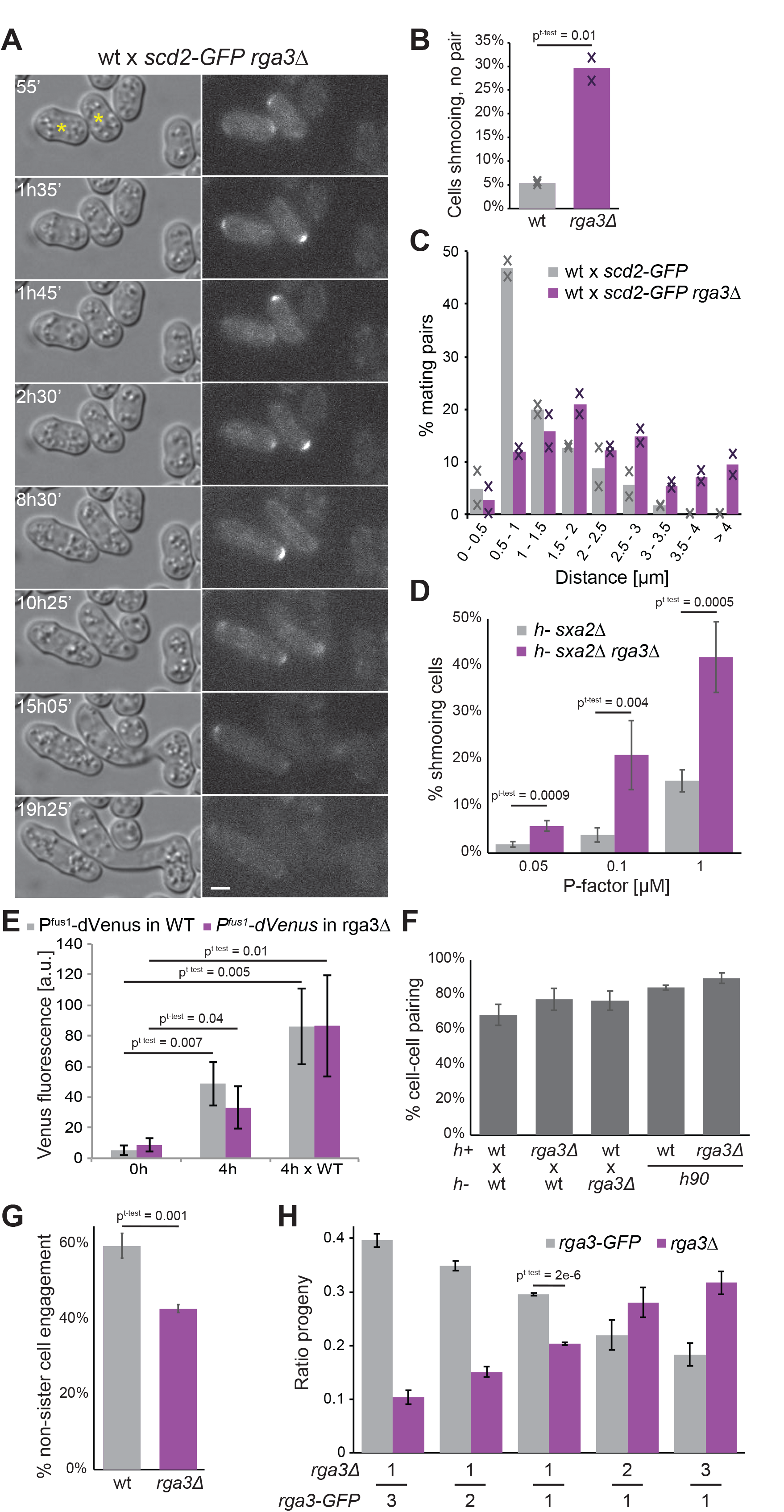
Rga3 limits growth projection in response to pheromone and confers a competitive advantage during sexual reproduction. (A) DIC and Scd2-GFP time-lapse images of h+ *rga3Δ scd2-GFP* cells mated to h-WT *scd2-mCherry* (not shown). Asterisks indicate *rga3Δ scd2-GFP* cells. Bar = 3μm. Time is in min from start of imaging. (B) Percentage of cells growing a cell projection that does not result in cell pairing in crosses of h+ WT or *rga3Δ* expressing Scd2-GFP with h-WT after 20 h on MSL-N pads. The average and individual data points from 2 experiments with n>1000 cells each are shown. (C) Quantification of the distance between partner cells prior to shmoo formation from mating mixtures as in A. n>60 in each of 2 experiments. (D) Percentage of h-*sxa2*Δ and h-*sxa2*Δ *rga3*Δ cells extending a growth projection 24h after 0.05, 0.1 or 1 μg/ml synthetic P-factor addition on MSL-N pads. n =4 experiments with > 1400 cells each. (E) Quantification of fluorescence of a double Venus transgene expressed under control of the *P^fus1^* promoter in cells before (0h) or after (4h) nitrogen starvation in absence or presence (x WT) of opposite mating type partners. N = 3 experiments. (F) Percentage of cell pair formation of heterothallic and homothallic wt and *rga3Δ* cells. n>1700 in 3 experiments (G) Percentage of zygotes formed by two non-sister cells in homothallic wt and *rga3Δ* cells. N = 3 experiments. (H) Quantification of the ratio of *rga3-GFP* and *rga3*Δ progenies resulting from a competitive mating assay, where an h-WT strain was mixed 1:1 with two h+ *rga3Δ* and h+ wt strains present in indicated ratio. N ≥ 3 experiments with n>170 colonies. Error bars show standard deviation.

The long shmoos of *rga3Δ* cells are reminiscent of pheromone hyper-signaling mutants (Bendezu and Martin, 2013). To test whether *rga3Δ* cells hyper-activate the pheromone-signaling pathway, we measured the fluorescence produced by Venus under control of the *P^fus1^* pheromone-responsive promoter (Petersen et al., 1995). Consistent with previous work, *P^fus1^*-driven Venus expression was induced upon sexual differentiation in nitrogen-starved cells in presence of mating partners. We also found that it was expressed, though at somewhat lower levels upon nitrogen starvation in absence of cells of opposite mating type. Expression levels were highly variable across the cell population, likely due to heterogeneity in the timing of sexual differentiation. However, in both cases, P^fus1^-Venus levels were indistinguishable in *rga3Δ* and WT cells, suggesting that the absence of Rga3 does not significantly increase the transcriptional output of the pheromone-signaling cascade, at least in this assay (Fig 4E). Similar observations were made using a *P^ste6^*-Venus reporter (data not shown).

We tested whether the formation of growth projections at lower pheromone concentrations in *rga3*Δ has physiological consequence on mating outcome. The overall mating efficiency, as measured as the percentage of cells engaged with a partner cell, was similar for *rga3*Δ and wildtype cells (Fig. 4F). However, we found that mate preference was significantly altered. Homothallic (self-fertile) wildtype cells can switch mating type such that two sister cells are potential mating partner (Klar, 2007), but preferentially mate with non-sister cells (Bendezu and Martin, 2013) (Fig. 4G). Conversely, homothallic *rga3Δ* cells displayed a small preference for sister cells (Fig. 4G). In competitive mating assays, *rga3+* cells also outperformed *rga3*Δ cells. We mixed different ratios of *rga3*Δ and *rga3-GFP* h+ cells labeled with distinct antibiotic markers (HPH- and G418-resistance, respectively) with wildtype h-cells marked with *myo52-tomato* (with NAT resistance) and quantified the proportion of HPH and G418-resistant spore progeny. *rga3Δ* cells were significantly depleted from the progeny of 1:1 mixes relative to *rga3-GFP* (Fig. 4H). This competitive disadvantage was also observed with 1:2 and 1:3 ratios of parental cells (Fig. 4H). We conclude that the more sensitive morphological response to pheromone in *rga3*Δ mutants decreases the ability of cells to efficiently pair with non-sister cells, producing a significant competitive disadvantage during sexual reproduction.

### Rga3 localizes to Cdc42 dynamic patches and promotes their exploratory dynamics

Similar to its localization during vegetative growth, Rga3 localized to sites of polarity during sexual reproduction. We labeled sites of polarity with the type V myosin Myo52. During early stages of the mating process, when Cdc42 undergoes dynamic polarization cycles, Rga3-GFP also co-localized with Myo52 (Fig. 5A), which was previously shown to label sites of Cdc42 activation (Bendezu and Martin, 2013). In shmooing cells, Rga3 localized to the shmoo tip, coincident with Myo52. At the site of cell fusion, Rga3 co-localized with the focus of Myo52 (Fig. S1C).

**Figure 5:**
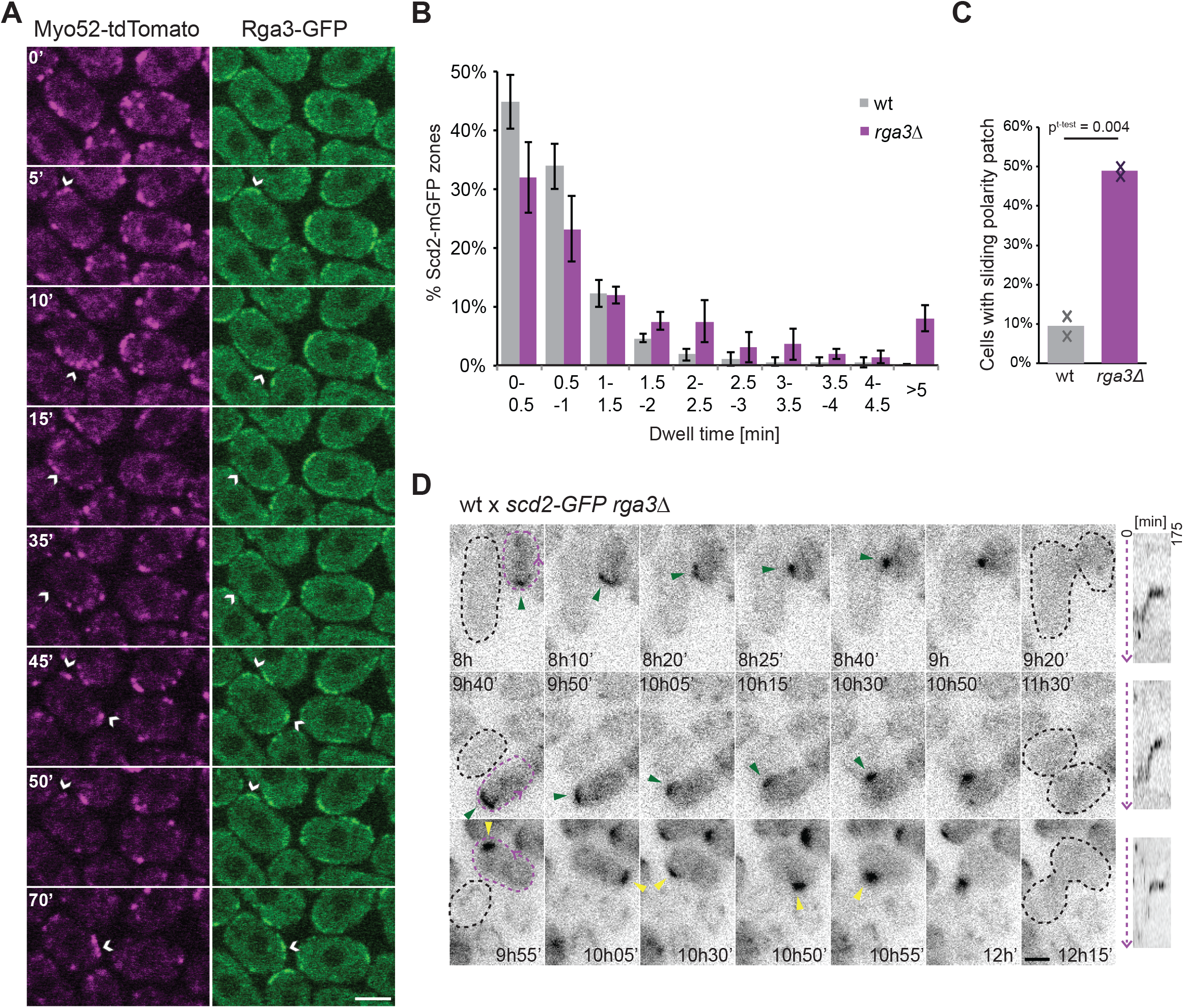
Rga3 localizes to Cdc42 dynamic patches and promotes their exploratory dynamics. (A) Medial spinning disk time-lapse confocal images of Rga3-GFP and Myo52-tdTomato in mating mixes during early stages of mating. Time is in minutes. Arrowheads highlight the coincidence of Rga3-GFP and Myo52-tdTomato. (B) Distribution of the dwell time of Scd2-GFP zones in h-WT and *rga3*Δ cells treated with 0.02 μg/ml synthetic P-factor on MSL-N pads. N = 4 experiments with n≥10 cells each. (C) Quantification of the frequency of sliding behavior for Scd2-GFP patches in h+ WT and *rga3*Δ cells mated with h-WT. n≥69 in each of 2 experiments. (D) Time-lapse images of Scd2-GFP in h+ *rga3*Δ cells (dashed purple) mated with h-WT (dashed black) showing two examples of Scd2-GFP patch sliding from the cell tip (top and middle) and one example of Scd2-GFP patch assembly-disassembly behavior prior to stabilization at the cell side (bottom). Green arrowheads indicate wandering patch movement. Yellow arrowheads highlight assembly-disassembly patch dynamics. To the right of the time-lapse are kymographs of *rga3*Δ cell cortex along the dashed purple arrow. All micrographs are medial plane spinning disk confocal images. Error bars show standard deviations. Time is in min from start of imaging. Bars = 3μm.

To relate the mating behaviors of *rga3Δ* to the activity state of Cdc42, we used Scd2-GFP as a marker for Cdc42-GTP and measured its dynamic properties upon pheromone exposure. In heterothallic *h-sxa2Δ* cells treated with 0.02μM P-factor, 90% of wildtype cells display an exploratory Scd2 patch with dwell time <1.5min (Bendezu and Martin, 2013; Merlini et al., 2016). Similarly-treated cells lacking *rga3* showed a more stable Scd2-GFP patch, with only 65% of cells displaying a dwell time <1.5min and nearly 10% of cells showing a non-dynamic patch, leading to cell shmooing (Fig. 5B), consistent with the higher pheromone sensitivity shown above (Fig 4D). In *rga3Δ* mating mixtures, consistent with the more persistent growth projections shown above (Fig 4B-C), the Scd2-GFP signal was often stable at the cell cortex for extended periods of time, with fewer cells exhibiting dynamic polarization. Notably, although *rga3Δ* cells like wildtype could engage partners from cell sides (26±7% and 29±5%, respectively), in 49% of these, the polarity patch first stabilized at the cell tip before exhibiting wandering motion to the cell side, rather than displaying the assembly-disassembly process typical of wildtype cells (Fig. 5C-D, Movies S2-S3). Similar patch wandering was observed in less than 10% of scd2-GFP wildtype cells. In sum, these data indicate that Rga3 modulates the dynamic behavior of the Cdc42 patch. Because Rga3 is recruited to polarity sites in response to Cdc42 activity, these data further suggest Rga3 may form a negative feedback promoting Cdc42-GTP zone turnover.

### Cells lacking all Cdc42 GAPs retain pheromone-dependent polarization ability

The effect of *rga3*Δ during mating incited us to investigate more globally whether Cdc42 GAPs contribute to the pheromone-dependent Cdc42 dynamic polarization. Unexpectedly, triple *rga3*Δ *rga4*Δ *rga6*Δ mutants retained ability to mate and fuse with wildtype cells (Fig. 6A). In addition, both Scd2-mcherry and CRIB-GFP still formed discrete zones that formed and disappeared around the cell periphery. Observation of Scd2 zones over long time periods revealed both instances of patch assembly-disassembly and wandering, as described above for *rga3*Δ (Fig. 6B, Movie S4). CRIB zones were often less well-defined than in wildtype cells, but also formed smaller zones than in vegetative cells (compare Fig. 6C to Fig. 2A). Upon cell pairing, both Scd2 and CRIB became clearly polarized near the mating partner towards which the triple mutant cells grew (Fig. 6B-C). We conclude that Cdc42 GAPs modulate the dynamic behavior of Cdc42-GTP patches but are dispensable for their assembly and disassembly. Thus, the formation of polarized zones of Cdc42-GTP relies on partly distinct regulatory mechanisms during the vegetative and sexual life cycles of the fission yeast.

**Figure 6:**
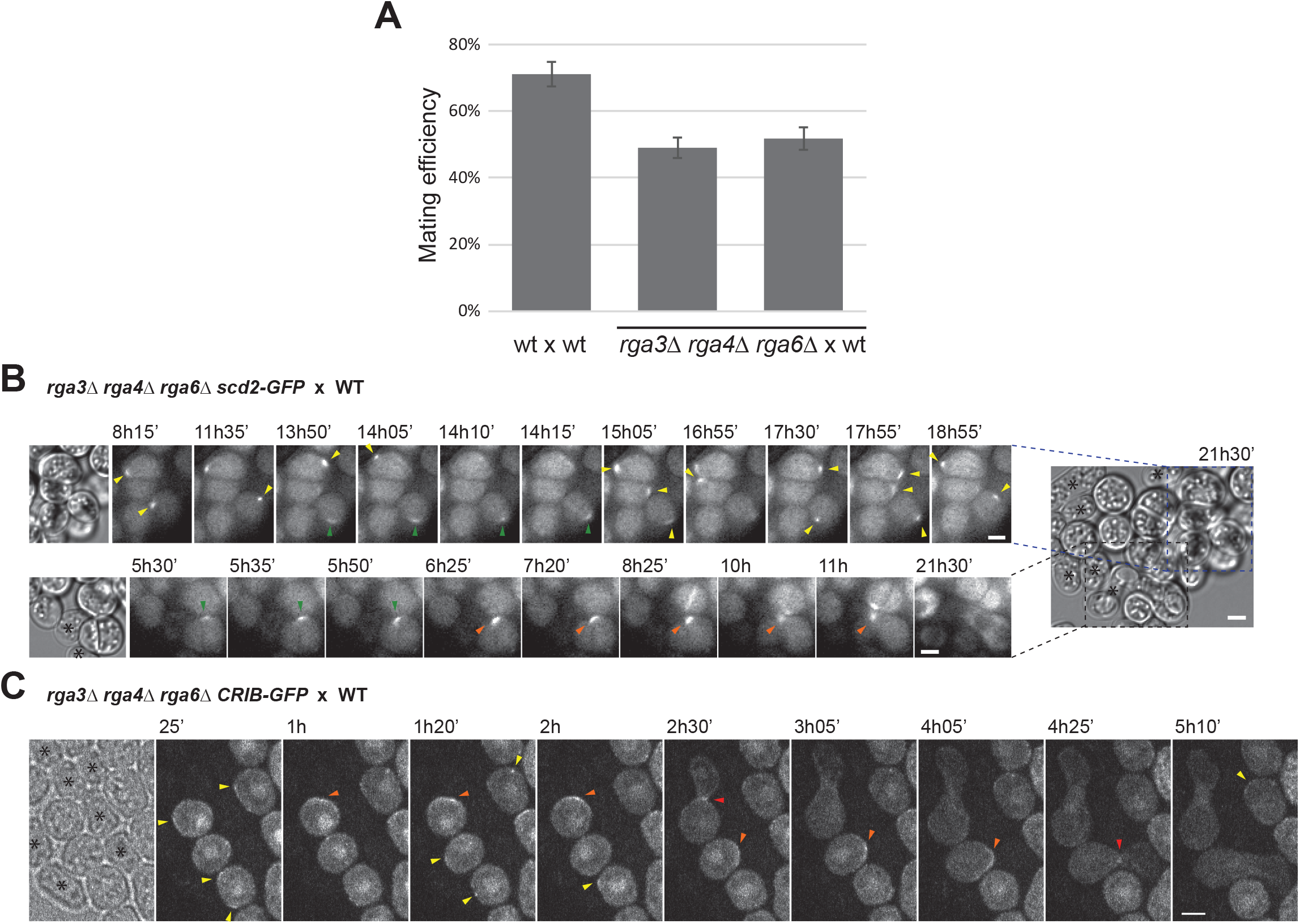
Cells lacking all three Cdc42 GAPs retain pheromone-dependent polarization ability. (A) Percentage of cell pair formation of heterothallic h-WT and *rga3*Δ *rga4*Δ *rga6*Δ cells (two independent strains) with h+ WT. n>1400 in 3 experiments. (B) Time-lapse images of Scd2-mCherry in h-*rga3Δ rga4Δ rga6Δ* cells mating with unlabeled h+ WT. Two examples extracted from the same large group of cells illustrate dynamic polarization (top) and shmoo formation (bottom). The first image is a DIC image on which the WT cells are marked with asterisks. Yellow arrowheads highlight assembly-disassembly patch dynamics. Green arrowheads indicate wandering patch movement. Orange arrowheads show the patch at the site of polarized growth. (C) Medial spinning disk confocal time-lapse CRIB-GFP images in h+ *rga3Δrga4Δrga6Δ* cells mating with unlabeled h-WT. The first image is a transmitted light image on which the WT cells are marked with asterisks. Yellow arrowheads highlight assembly-disassembly patch dynamics. Orange arrowheads indicate the presence of the patch at the site of polarization. Red arrowheads point to fusion with partner cells. Time is in min from start of imaging.

## DISCUSSION

Spatial regulation of Cdc42 GTPase activity is critical for cell polarization. In this study, we identified and studied the function of Rga3, a novel Cdc42 GTPase activating protein. Bioinformatics analysis revealed that Rga3 is a paralog of the previously characterized Rga4, with strong sequence homology throughout their sequences, except for the unique presence in Rga3 of a C1 domain, a lipid-binding domain that typically binds phorbol esther and diacyl glycerol (Das and Rahman, 2014). The Rga3 domain architecture is found in related GAPs in basidiomycetes, as well as in the *Taphrinomycotina* group, a basal ascomycete lineage that includes the fission yeast clade and *Pneumocystis* pathogens. The association of C1 and RhoGAP domain is also present in other eukaryotic lineages including metazoans, in the Chimaerin family of RacGAPs (Yang and Kazanietz 2007), which contains the GAP domain most closely related to that of Rga3 and Rga4. By contrast, GAP proteins in other ascomycete species lack C1 domains altogether. These observations suggest that the association of C1 and RhoGAP domains represents the ancestral state of this gene family. A possible evolutionary scenario is that during early ascomycete evolution, an ancestral C1-containing GAP gene underwent duplication with one gene copy subsequently loosing its C1 domain. The fission yeast clade retained both gene copies, yielding Rga3 and Rga4, whereas one copy was lost in all other ascomycetes.

Duplication of the ancestral gene must have been accompanied by evolution of alternate membrane localization determinants in the two resulting genes. Indeed, Rga3 and Rga4 decorate distinct, complementary cortical regions. In Rga3, the C1 domain is necessary and sufficient for plasma membrane localization, consistent with its predicted lipid-binding activity, but does not confer on its own specificity for cell poles. Localization to sites of polarity also requires the LIM domains and coiled-coil regions of Rga3, likely through protein-protein interaction, but interestingly not the GAP domain. Of note, none of the Rga3 fragments tested displayed Rga4-like cell side localization, suggesting that localization to sites of Cdc42 activity may represent the ancestral state. Consistent with this idea, the related *S. cerevisiae* GAPs also localize to sites of polarity (Caviston et al., 2003). Whether the extant cell side-localization of Rga4 masks an ancestral polar localization remains to be tested. Interestingly, the observation that Rga3 is recruited by GTP-locked Cdc42, yet its localization at sites of polarity is independent from its GAP domain, suggests this recruitment does not simply occur through direct binding of the GAP domain to Cdc42-GTP but indirectly, for instance through protein-protein interaction with a Cdc42-GTP effector. In this way, Rga3 may form a negative feedback on Cdc42 activity.

### Rga3 promotes Cdc42-GTP zone instability during mating

The maintenance of duplicated gene copies in the fission yeast genome and the unique localization of Rga3 at the polarity patch suggest Rga3 (or Rga4) acquired a specific function, distinct from that of the two other GAPs. During mitotic growth however, though Rga3 participates in Cdc42 inactivation with Rga4 and Rga6, it does not appear to play any unique function. Even analysis of Cdc42-GTP oscillatory patterns did not show crucial changes in *rga3Δ* mutants, with only a reduction in the oscillation frequency observed, difficult to reconcile with any direct role for a negative regulator. By contrast during mating, the absence of Rga3 led to noticeable changes in Cdc42 patch dynamics: the Cdc42 patch exhibited a longer dwell time leading to exacerbated outgrowth and showed reduction in assembly-disassembly behavior.

One question is whether Rga3 directly promotes patch destabilization or acts more indirectly by lowering downstream signaling. Previous work indeed showed that patch stabilization occurs upon increased signaling in strains lacking negative pheromone-GPCR control (Bendezu and Martin, 2013), which signals through a MAPK cascade proposed to be partly under Cdc42 regulation (Marcus et al., 1995; Tu et al., 1997; Weston et al., 2013). However, we could not detect significant increase in transcriptional reporters of MAPK activity in *rga3Δ* cells. This suggests that MAPK output is not significantly altered in absence of Rga3, though we note that small changes are unlikely to be detected with this assay, due to high cell-to-cell variability. Thus, we favor the view that, by being recruited to the polarity patch, Rga3 directly promotes patch destabilization by reducing Cdc42 activity.

Remarkably, the cell is largely able to compensate the reduction in assembly-disassembly behavior upon loss of Rga3 through an alternate patch sliding strategies. Indeed, although *rga3Δ* cells exhibit measurable disadvantage when placed in competitive mating situation, they remain able to extend a growth projection from their side and display unaltered mating efficiency when unchallenged. Patch sliding is normally rarely observed in *S. pombe*, but it is the principal mode of gradient tracking in *S. cerevisiae*. In this case, movement of the patch up-gradient is proposed to rely on dilution effects from vesicle delivery and on feedback regulation through local signal of the pheromone GPCR-MAPK (Dyer et al., 2012; Hegemann et al., 2015; McClure et al., 2015). The patch wandering behavior observed in *rga3Δ* cells may rely on similar mechanisms, which may exist also in wildtype fission yeast cells to refine the position of the patch. It is interesting that the presence or absence of a single Cdc42 GAP partly converts patch dynamic behavior from assembly-disassembly to wandering, suggesting that cells may naturally also be able to switch between these two modes of cell pairing.

### Different GAP requirement for Cdc42-GTP polarization in distinct life stages

One surprising observation is the very distinct cell polarization outcome of the absence of the three Cdc42 GAPs Rga3, Rga4 and Rga6 during the mitotic and sexual life cycles. Mitotic cells lacking the three GAPs formed very large Cdc42-GTP domains that could extend over more than half of the cell surface, and therefore acquired a round cell shape. This phenotype, similar to expression of GTP-locked Cdc42, is consistent with the idea that spatial restriction of Cdc42 activity is a dynamic process that relies not only on local activation at cell tips, but also on GTP hydrolysis to counteract Cdc42 diffusion. An essentially identical outcome is observed for Ras1 GTPase normally selectively active at cell poles: upon deletion of Ras1 only GAP, the GTPase is now active around the entire cell cortex (Merlini et al., 2018). By contrast, cells lacking all three GAPs were still almost fully mating-competent and retained the ability to polarize active Cdc42, as labeled by the CRIB reporter or the Cdc42-GTP-binding Scd2 scaffold. We conclude that the three Cdc42 GAPs studied here, though modulating patch dynamic properties, are not essential for Cdc42-GTP polarization during mating and that the polarization mechanisms of Cdc42-GTP can be distinct even in different life stages of the same cell. This finding represents an interesting paradigm of the emerging theme that polarity mechanisms show important context-dependent variation (St Johnston, 2018).

A key question is what accounts for this difference. During mitotic growth, cells polarize in response to cell-intrinsic landmarks deposited at cell poles by microtubules (Martin, 2009). As microtubules orient along cell length by sliding along cell sides (Martin and Arkowitz, 2014), the loss of cell shape in triple GAP mutants likely does not permit landmark positioning, in turn compounding the effect of the GAP deletions. By contrast during mating, cells polarize in response to extrinsic pheromone gradients (Merlini et al., 2013). Thus, one possibility is that the positional cue provided by external pheromone gradients is sufficient for polarization even from a round cell shape. However, this is likely a too simplistic view, as GAP-deleted cells without close WT partners, unlikely to experience any sharp pheromone gradients, are also seen to efficiently polarize, often in direction unrelated to that of candidate partner cells (see Fig. 6B). It is possible that there exists yet another Cdc42 GAP specific for the sexual life cycle. However, none of the other encoded GAP genes appear to increase levels upon pheromone exposure (Mata and Bahler, 2006), though this does not exclude other post-transcriptional regulations. Alternatively, Cdc42-GTP polarization may depend on other modes of negative regulation during mating. Possibilities include GDI-mediated membrane extraction, which plays little role during mitotic growth (Bendezu et al., 2015; Nakano et al., 2003); endocytosis-dependent Cdc42-GTP retrieval from the membrane (Orlando et al., 2011; Slaughter et al., 2009; Watson et al., 2014); or a GAP-independent boost of the intrinsic Cdc42 GTPase activity promoting the return of Cdc42 to the basal state. Indeed, the intrinsic GTPase activity of Cdc42 we observed in vitro (Fig. 2C; Cdc42-GTP t_1/2_ < 5 min) is relatively high compared to that of Cdc42 in other species (Zhang et al., 1999) or Ras1 in *S. pombe* (Merlini et al., 2018), and may thus suffice to be effective against an estimated lateral diffusion of Cdc42-GTP of 2μm^2^/min (Bendezu et al., 2015).

## MATERIALS AND METHODS

### Strain construction, media and growth conditions

Strains used in this study are listed in Table S1. All plasmids used are listed in Table S2. Standard genetic manipulation methods for *S. pombe* transformation and tetrad dissection were used. For microscopy of fission yeast cells during exponential growth, cells were grown in Edinburgh minimal medium (EMM) supplemented with amino acids as required. For biochemistry experiments, cells were grown in rich Yeast extract medium (YE). For assessing cells during the mating process, liquid or agar minimal sporulation medium without nitrogen (MSL-N) was used (Egel et al., 1994; Vjestica et al., 2016). All live-cell imaging during sexual lifecycle was performed on MSL-N agarose pads (Vjestica et al., 2016). Mating assays were performed as in (Vjestica et al., 2016).

All cells were grown at 25°C, unless differently specified. For cell length measurements all strains had identical auxotrophies and were grown in EMM-ALU to mid-log before Calcofluor (Sigma) addition to a final concentration of 5μg/ml from a 200X stock solution. For plasmid expression, all strains were grown in EMM-AU for 20 hours before imaging, unless differently specified.

Construction of YSM3149 in Fig. 3F was obtained by integration of the C-terminal part of the gene, lacking the C1 domain, at its genomic locus. First, an expression plasmid was generated: Rga3 GAP domain was amplified from pSM1337 (see table S2) with primers osm4400 (5’-CGggatccGGGTCGGTTTTGCCTCAAGTAATTG) and osm4401 (5’-CCCcccgggGCCAGATTATGAAGAATCTTATCCA), digested with XmaI and BamHI and ligated to similarly treated pSM1339 (see table S2) to generate plasmid pREP41-rga3(-C1)-eGFPc. Second, 819bp from the C-terminal part of this plasmid was amplified with osm4770 (5’-CGACGGATCCCCGGGTTAATtaaATTCAGCATAGAGTACC) - osm4769 (5’-TTCTTCTCCTTTACTGTTAattaaCAGATTATGAAGAATC) and cloned with the In-Fusion cloning kit (Takara) into pSM206 (pFA6a-GFP(S65T)-kanMX6) previously digested with PacI, resulting in pSM2197. Finally, standard *S. pombe* transformation was performed on YSM3130 by amplification of pSM2197 fragment with primers osm774 (5’-GATTTACAGCATGCTACCAAAAAGAGTACGGCTCTTCAGTTTATGCTTGATAACGTGGATAAGATTCTTCATAATCTGCGGATCCCCGGGTTAATTAA) – osm773 (5’-TTATTATTATCGTAAAGTTCCTTCTTCCCTAGAAAAATACAAGTTCCCCGATATTTTAATCATTATTCATAAAACAAGGAATTCGAGCTCGTTTAAAC).

YSM3156 and YSM3157 were obtained by cross with a strain previously transformed with pSM2187 (see Table S2).

### Mating Assays

Mating assays were performed as in (Egel et al., 1994). Briefly, pre-cultures of cells were grown over night in MSL+N at 25°C to reach an OD_600_ of 0.5-1. Cultures were then diluted to an OD_600_ of 0.025 in MSL+N and grown for 16-20 hours to an OD_600_ of 0.5-1 at 30°C. Cells were washed three times with MSL-N, diluted to an OD_600_ of 1.5 in 1-3 ml MSL-N and incubated at 30°C for 1-4 h (depending on the mating stage to be visualized). Cells were mounted onto MSL-N agarose pads (2% agarose) before imaging in overnight movies at room temperature (about 22°C) or incubated at 18°C overnight before imaging. Mating efficiency was measured as in (Dudin et al., 2015; Dudin et al., 2016). Sister cell engagement and zone dwell time were measured as in (Bendezu and Martin, 2013).

For pheromone treatments, P-factor pheromone was purchased from Pepnome (Zhuhai City, China) and used from a stock solution of 1 mg/ml in methanol. Different concentrations of pheromones were directly added to the melted MSL-N agarose before mounting cells on the pads. Cells were then imaged overnight or incubated at 18°C overnight before imaging.

For Fig. 4D, the pheromone treatment was performed on pads and left at 25°C for 24 hours.

### Competition assay

Mating assays were performed by mixing YSM1232 (h+), YSM3160 (h+) and YSM952 (h-) (see table S1) at different ratios. Cell mixes were plated on MSL-N plates for 24 hours at 25°C. The cells were then transferred into an Eppendorf tube and treated with glusulase (1:200 from stock solution; PerkinElmer) in H_2_O overnight at 25°C. Only spores survived the treatment as confirmed by microscopic inspection. Spores were then plated on YE and grown for 3 days at 30°C. The plates were then replicated on selective media in order to determine the genotype of each colony.

### Microscopy

Microscopy was performed on live cells, either on a spinning disk confocal microscope or on a DeltaVision epifluorescence system. Spinning disk microscopy was carried out using a Leica DMI4000B inverted microscope equipped with HCX PL APO X100/1.46 (NA) oil objective and a PerkinElmer Ultraview Confocal system (including a Yokagawa CSU22 real-time confocal scanning head, and solid-state laser and a cooled 14-bit frame transfer EMCCD C9100-50 camera).

For cell-length measurements, images of calcofluor-stained cells were taken on a DeltaVision platform (Applied Precision) composed of a customized Olympus IX-71 inverted microscope and a UPlan Apo 100X/1.4 NA oil objective, a CoolSNAP HQ2 camera (Photometrics), and an Insight SSI 7 color combined unit illuminator.

#### Image Analysis

For cell length and width measurements on calcofluor stained cells, a line was drawn manually across the length and width of septated cells from the middle of one tip to the other and the length measured using the ImageJ Measure tool.

#### Fluorescence image quantification

Quantification of Cdc42-mCherry^SW^ and CRIB-GFP distribution in Fig. 2A was done on the sum projection of five consecutive images. The intensity profile along a 3 pixel-wide segmented line along half of the cell cortex was measured for at least 20 tips for both red and green channels. Profiles were centered on the maximum pixel value for the CRIB channel and subsequently split into two half tips.

Quantification of Cdc42-mCherry^SW^ and Rga3-GFP distribution in Fig. 3B was done on the sum projection of 5 consecutive images. The intensity of a 3 pixel wide segment was collected from 24 half-cell tips for both red and green channels. The values were normalized to the maximum and minimum pixel values.

Quantification of Cdc42-mCherry^SW^ (wildtype and Q61L, expressed from plasmid) and Rga3-GFP distribution in Fig. 3C was done on the sum projection of ten images obtained collected every 30 seconds for 5 minutes. The intensity of a 3 pixel wide segment was collected from 18 half-cell tips for both red and green channels. Only rod-shaped cells were taken into analysis.

Quantification of Rga3-GFP (wildtype and ΔC1) distribution in Fig. 3F was done on the sum projection of five consecutive images. The intensity of a 3 pixel wide segment was collected from at least 20 tips for the green channel. Total intensity quantification was performed by drawing a line along the cell cortex with ImageJ polygon tool and subtracting the background.

For Fig. 4E, wildtype or *rga3Δ* h+ cells expressing a *P^fus1^-double-Venus* reporter were treated as for mating assays. After shift on MSL-N cells were mixed with wildtype cells expressing Scd2-mCherry that were either h+ (for analysis in absence of mating partners) or h- (for analysis in presence of mating partners). Scd2-mCherry cells were used to measure background fluorescence in the Venus channel. Samples were imaged just after mixing (time 0) or after 4h. Cell fluorescence was measured in ImageJ by placing a square in the middle of each cell and calculated as: (P^fus1^-Venus signal – local background)^t=×^ – average of (Venus signal in Scd2-mcherry cells – local background)^t=x^. Average of at least 20 cells per each timepoint was calculated.

Kymographs in Fig. 5D were constructed in ImageJ version 1.51 (NIH) by drawing a 3-pixel-wide line around the cell cortex and using the ImageJ Reslice tool.

#### CRIB oscillations

The analysis on CRIB-GFP oscillatory behavior was done as in (Das et al., 2012). Briefly, time-lapse movies of wild-type and *rga3Δ* cells expressing CRIB-GFP were obtained. Cells were imaged on EMM-ALU 2% agarose pads every 1 minute for 30 minutes by spinning disk confocal microscopy. 20 cells were used for the analysis. Only bipolar cells were taken into consideration. CRIB-GFP intensity was obtained by drawing a 3 pixel wide segment at each cell tip. The traces were normalized to a linear regression of another cell intensity over time to compensate for fluorescence loss due to imaging, normalized to the maximum value and plotted against time. The frequency was calculated by identifying individual peaks on the plot and calculating the average time interval between them. Cross-correlation was calculated using the Excel Correl function.

### Biochemistry methods

#### Recombinant protein production

Expression of GST-proteins or GST was induced in BL21 bacteria from pGEX-4T-1–derived plasmids. In brief, cells were grown overnight in LB (Luria–Bertani) medium supplemented with 100 μg/ml ampicillin at 37°C. 250 ml LB-ampicillin were inoculated with 6.25 ml of the saturated culture and grown for 3 h at 37°C. Protein expression was induced by the addition of 100 μM IPTG for 5 h at 18°C. Bacterial pellets were resuspended in 5 ml PBS (137mM NaCl, 2.7mM KCl, 1.4mM KH2PO4, 4.3mM NaH2PO4, pH 7.4), digested with 1 mg/ml lysozyme, treated with 1 μg/ml DNase I, sonicated 6x for 30 s (40% amplitude), incubated with 1% Triton X-100 in PBS buffer at 4°C for 30 min, and centrifuged 15 min at 4°C at 10,000 g. Soluble extract was incubated with 200 μl glutathione–Sepharose beads at 50% slurry for 2 h at 4°C. Finally, beads were washed 3x with cold PBS and eluted in four steps in 100 μl elution buffer (15 mM reduced glutathione, 50 mM Tris-HCl, pH 8).

Protein concentration was determined from SDS-PAGE gels by using bovine serum albumin (BSA) as a standard protein at different concentrations (4, 2, 0.8, 0.4 and 0.2 μg/μl).

#### Cdc42-GTP pull-down

The Cdc42-GTP pull-down assay was performed as in (Tatebe et al., 2008). Cdc42-mCherry^SW^ strains grown overnight at 30°C in 150ml YE media were used to produce protein extracts as in Revilla-Guarinos et al., 2016. Briefly, cell extracts from wild-type, *rga3Δ, rga4Δ, rga6Δ* and *rga3Δrga4Δrga6Δ* cells expressing Cdc42-mCherry^SW^ were obtained by mechanical breakage of the cells using glass beads and FastPrep. Cells were resuspended in 500 μl of lysis buffer (50 mM Tris-HCl, pH 7.5, 20 mM NaCl, 0.5% NP-40, 10% glycerol; 0,1 mM DTT, 2 mM MgCl2). Protein extracts were then incubated with bacterially expressed GST-CRIB (see “Recombinant protein production” section) and sepharose slurry, for 1 hour at 4°C in binding buffer (25 mM Tris-HCl, pH 7.5, 1 mM DTT, 30 mM MgCl2, 40 mM NaCl, 0.5% Nonidet P-40). The bead pellet was then washed 3 times with 25 mM Tris-HCl, pH 7.5, 1 mM DTT, 30 mM MgCl2, 40 mM NaCl, 1% Nonidet P-40 and 2 times with the same buffer without Nonidet P-40.

Samples were loaded on 12% protein gel and resolved by SDS-PAGE for Western blot analysis. Standard protocols were used for SDS-PAGE and Western blot analysis. Antibodies used on western blots were: mouse monoclonal [6G6] to RFP (Chromotek).

#### GAP assay

1 μg of purified Cdc42 (see “Recombinant protein production” section) was incubated in 70 μl of loading buffer (20 mMTris-HCl, pH 7.5, 25 mM, NaCl, 5 mMMgCl2, and 0.1 mM DTT) containing 1.5 μl [γ-32P]GTP for 10 min at 30 °C. The mixture was then put on ice, and 10 μl were added to 50 μl of reaction buffer (20 mM Tris, pH 7.5, 2 mMGTP, and 0.6 μg/μl BSA) with 5 μg of GAPs (see “Recombinant protein production” section) or 50mM TRIS pH8. The tube was transferred at 30 °C, and every 2 minutes, 10 μl of the reaction was diluted in 990 μl cold washing buffer (20 mMTris-HCl, pH 7.5, 50 mM NaCl, and 5 mM MgCl2). The samples were filtered through pre-wetted nitrocellulose (Millipore) filters, washed three times with 4 ml cold washing buffer and air-dried. The amount of filter-bound radioactive nucleotide was determined by scintillation counting.

### Sequence Analysis

All RhoGAP-containing proteins on the Pfam site (on 12th June 2017) from the following species were selected: Ascomycetes: *A. nidulans; P. rubens; A. oryzae; C. posadasii; P. tritici-repentis; N. crassa; T. deformans; S. pombe; S. japonicus; S. cryophilus; P. murina; P. jirovecii; P. pastoris; K. lactis; S. cerevisiae; A. gossypii; Z. rouxii; Y. lipolytica;* Basidiomycetes*: U. maydis; S. reilianum; R. toruloides; M. larici-populina; T. mesenterica; C. neoformans*, and sequences of the RhoGAP domain aligned using the Clustal W method in MegAlign (Lasergene). This identified Rga3 as an Rga4 paralog. An alignment using whole protein sequences from entries on the same phylogenetic branch as Rga3 and Rga4 was subsequently performed and is shown in Fig. 1C. Domain architecture was derived from SMART analysis.

Figures were prepared with ImageJ64 and assembled using Adobe Illustrator CS5. All error bars are standard deviations. All experiments were done at least three independent times, except for those described in figures 4B, 4C, 5C.

**Table S1:**
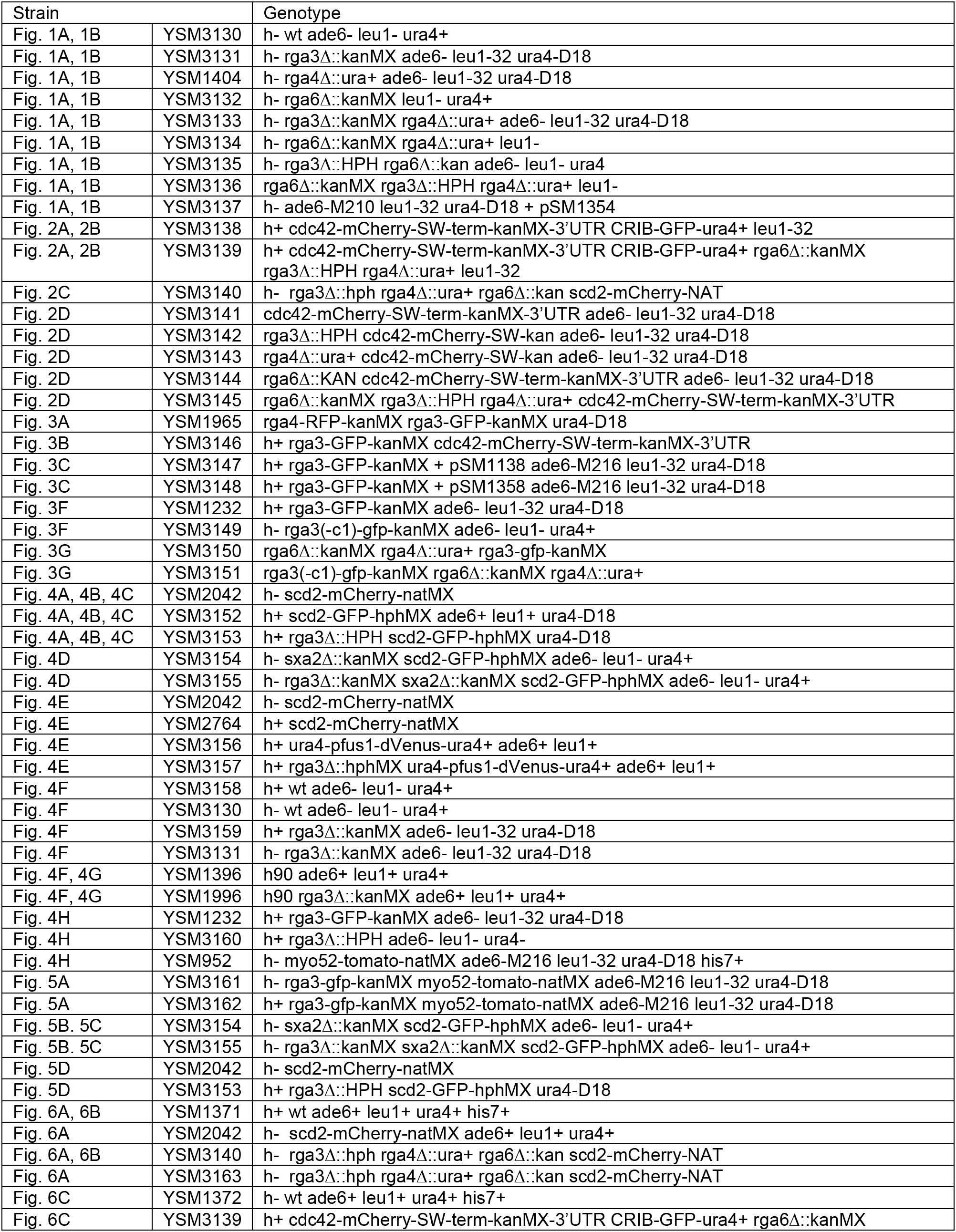
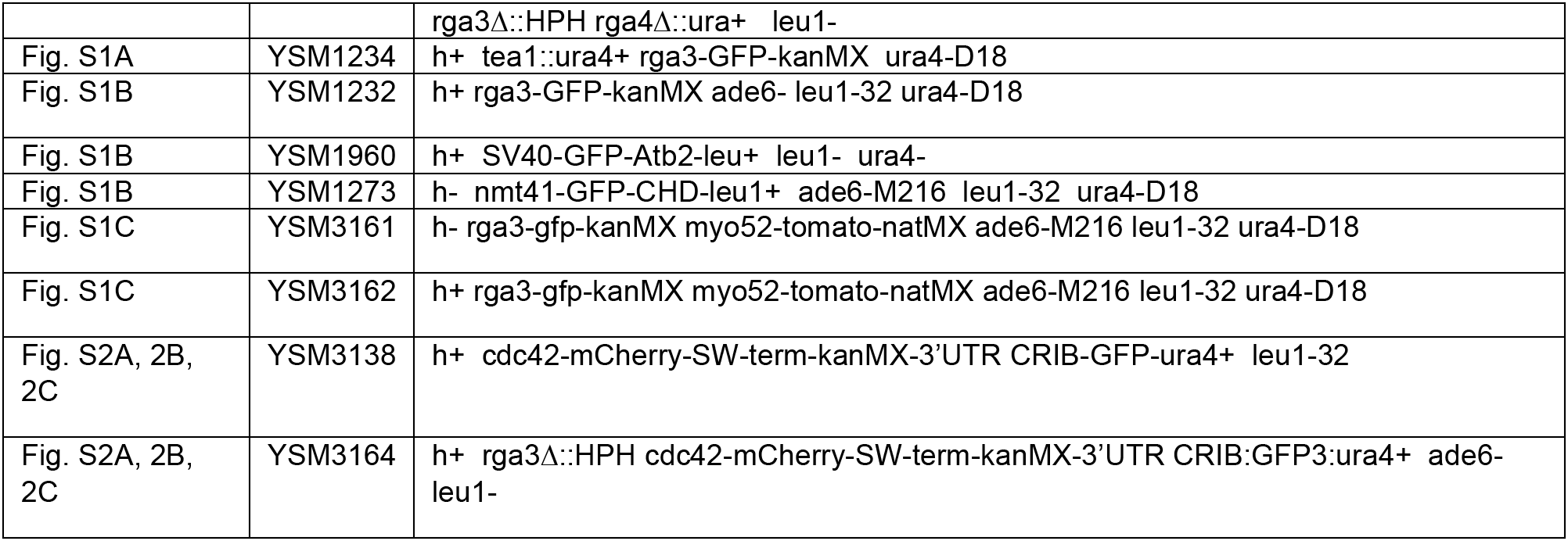
Strains used in this study

**Table S2:**
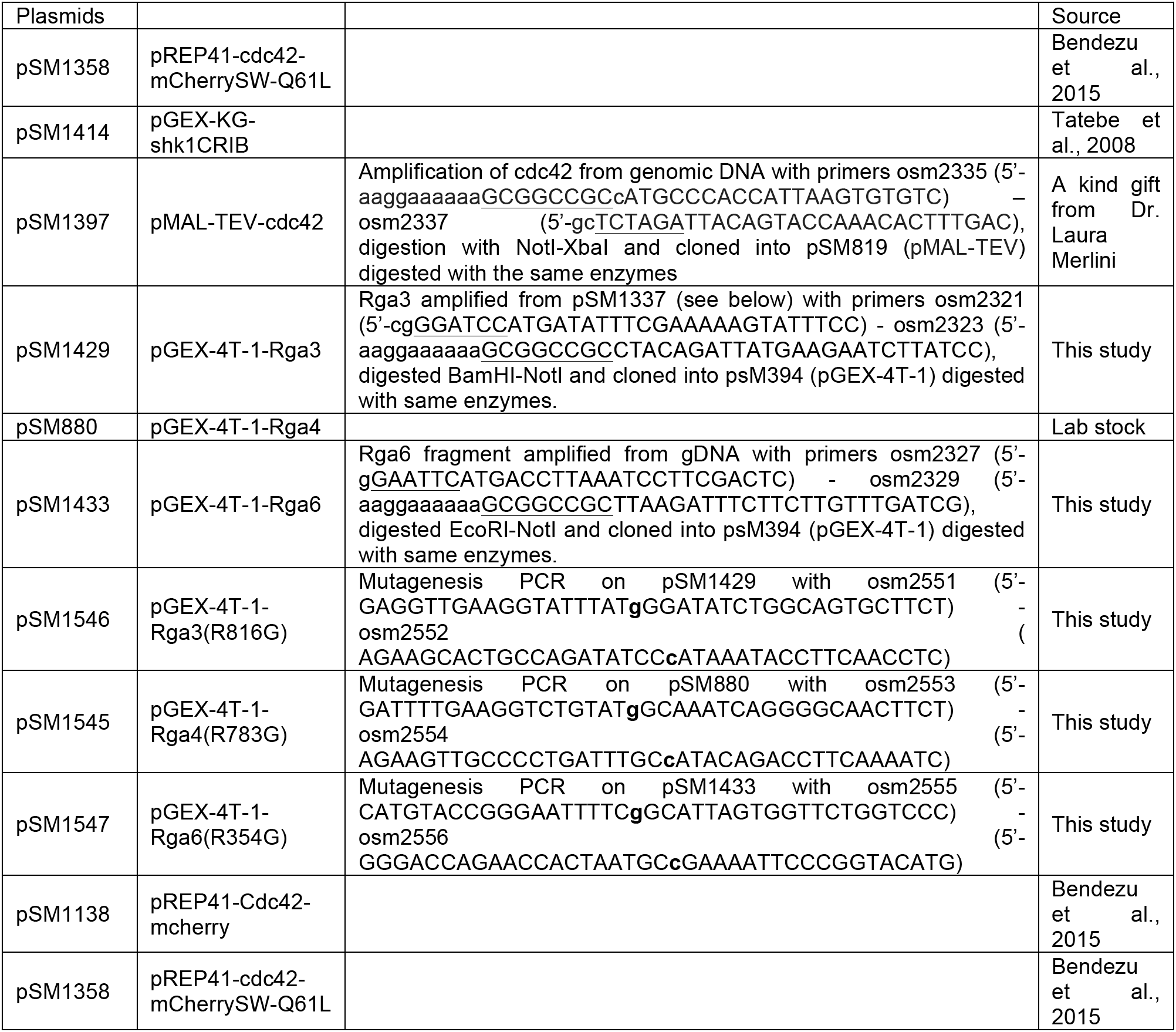
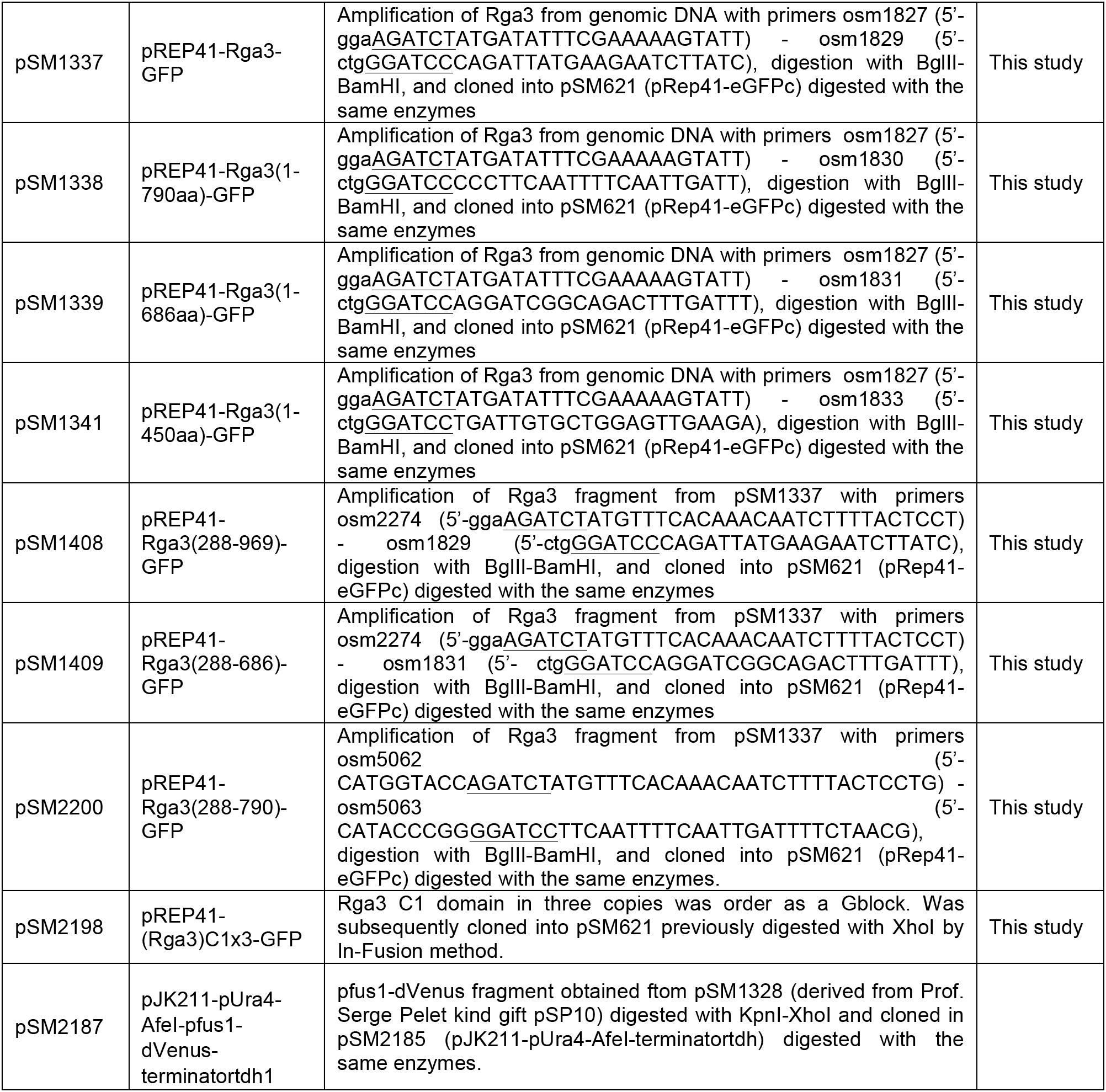
Plasmids used in this study. Restriction enzymes are underlined; point mutagenesis are bold.

C1×3 GBlock sequence:

5’

tacc<u>AGATCTCTCGA</u>GTGTTTCACGTTAATGCAATATTCAAGCCCTCAAGGTGTTATATTTGCT
CGGAGAGCGTATGGGGATCTGAACTCCGCTGCTTCCATTGCTCAATCTCATGCCATTCGC
GGTGTTTAAAAAGGCTGTTTGCCGAGTCGGTTTTCCATGTCAATGCCATCTTCAAACCTTCC
CGCTGCTATATATGCTCTGAAAGTGTTTGGGGTTCTGAGTTACGTTGTTTCCACTGTTCTAT
TTCGTGTCATTCCAGATGCCTTAAGCGTTTGTTCGCTGAGTCCGTCTTTCACGTTAACGCTA
TTTTCAAGCCCTCTCGATGTTACATTTGTTCCGAATCGGTATGGGGCTCTGAACTTAGGTG
CTTCCATTGTTCCATTTCTTGCCACTCTCGCTGCTTGAAGAGATTATTTGCCGAGTCTGGAT
CCccgggtatga

Manual codon degeneration was required to avoid sequence repetitions.

---

**Figure S1.**
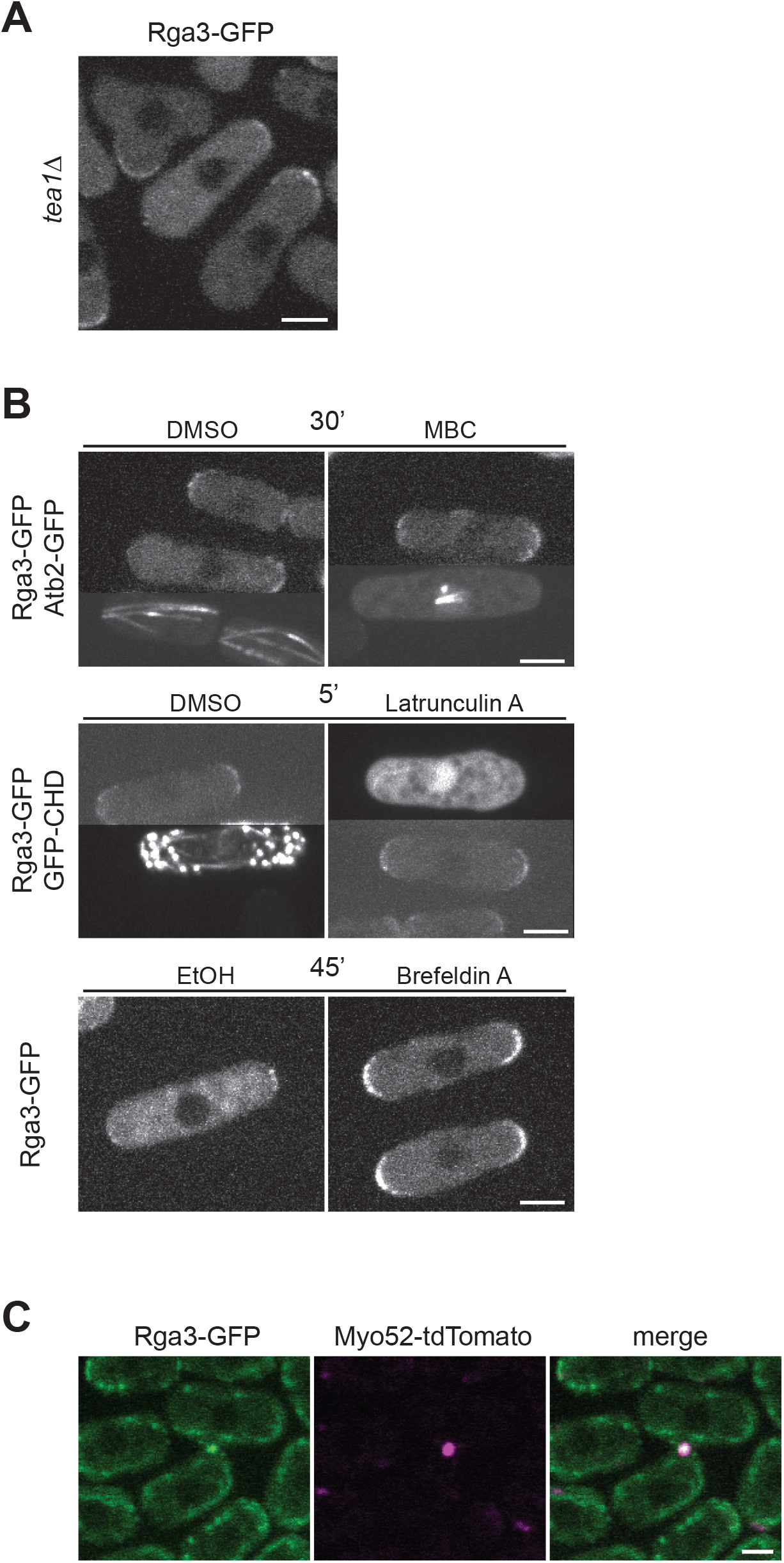
Localization of Rga3 to sites of polarity independent of the cytoskeleton. (A) Rga3-GFP at the single growing cell pole of tea1Δ cells. (B) Rga3-GFP in cells treated with MBC (top), Latrunculin A (middle) or Brefeldin A (bottom). Efficiency of MBC and LatA was confirmed by co-imaging cells carrying GFP-labeled Atb2 (alpha-tubulin) or GFP-CHD, which labels F-actin, respectively. Images are contrasted differently for these markers. (C) Localization of Rga3-GFP (green) at the site of cell-cell fusion, as labeled by the Myo52 fusion focus (purple). Bars are 3μm.

**Figure S2.**
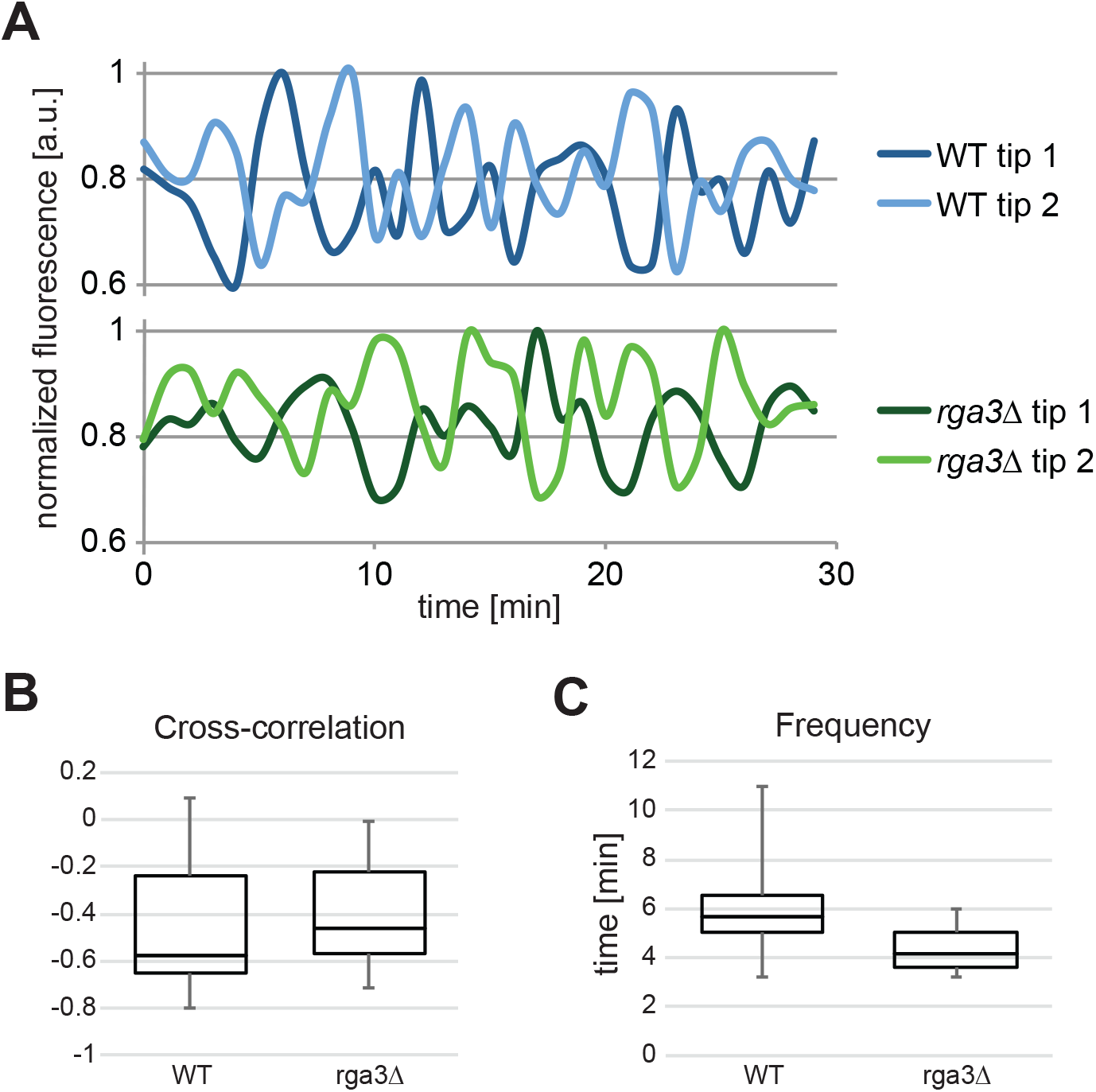
Minor changes in Cdc42-GTP oscillations in *rga3*Δ cells. (A) Relative fluorescence intensity of CRIB-GFP at the two cell poles of a single WT and *rga3Δ* cell over 30 min. (B) Values for cross-correlation between the fluorescence intensities at the two cell poles in cells as in A, showing anti-correlation in both WT and *rga3Δ*. (C) Frequency of oscillation derived from traces as in A. The graphs show median values, and upper and lower quartiles. Error bar indicate max and min values. n = 20 cells.

**Movies S1. Exacerbated growth phenotypes of *rga3Δ* cells during mating.**

DIC and Scd2-GFP time-lapse images of h+ *rga3Δ scd2-GFP* cells mated to h-WT. From the two *rga3Δ scd2-GFP* cells in the middle of the image, the top one extends a long shmoo and mates with a WT cell while the bottom polarizes growth predominantly at the top cell pole, but intermittently at the bottom one, without finding a partner cell. Time is in min. See also Fig. 4A.

**Movie S2. Example of *rga3Δ* cells with patch wandering behavior.**

Time-lapse images of Scd2-GFP in h+ *rga3Δ* cells mated with h-WT. Two rga3Δ scd2-GFP cells, marked with asterisks, form patches at cell poles that then “slide” towards the mating point with the partner cell. Time is in min.

**Movie S3. Example of *rga3*Δ cells with patch assembly-disassembly behavior.**

Time-lapse images of Scd2-GFP in h+ *rga3Δ* cells mated with h-WT. Example of an rga3Δ scd2-GFP cell (marked with asterisk) that assemble and disassemble cortical patches before mating with a partner cell on the side. Time is in min. See also Fig. 5D, bottom cell.

**Movie S4. Cells lacking Rga3, Rga4 and Rga6 retain polarization ability during mating.**

Time-lapse images of Scd2-mCherry in h-*rga3Δ rga4Δ rga6Δ* cells mating with unlabeled h+ WT. See also Fig 6B. Time is in min.

